# Global population divergence and admixture of the brown rat (*Rattus norvegicus*)

**DOI:** 10.1101/065458

**Authors:** Emily E. Puckett, Jane Park, Matthew Combs, Michael J. Blum, Juliet E. Bryant, Adalgisa Caccone, Federico Costa, Eva E. Deinum, Alexandra Esther, Chelsea G. Himsworth, Peter D. Keightley, Albert Ko, Åke Lundkvist, Lorraine M. McElhinney, Serge Morand, Judith Robins, James Russell, Tanja M. Strand, Olga Suarez, Lisa Yon, Jason Munshi-South

## Abstract

Once restricted to northern China and Mongolia, the brown rat (*Rattus norvegicus*) now enjoys a worldwide distribution due to the evolution of commensalism with humans. In contrast to black rats and the house mouse, which have tracked the regional and global development of human agricultural settlements, brown rats do not appear in the European historical record until the 1500s, suggesting their range expansion was a response to relatively recent increases in global trade and modern sea-faring. We inferred the global phylogeography of brown rats using 32k SNPs to reconstruct invasion routes from estimates of population divergence and admixture. Globally, we detected 13 evolutionary clusters within five expansion routes. One cluster arose following a southward expansion into Southeast Asia. Three additional clusters arose from two independent eastward expansions: one expansion from Russia to the Aleutian Archipelago, and a second to western North America. Rapid westward expansion resulted in the colonization of Europe from which subsequent colonization of Africa, the Americas, and Australasia occurred, and multiple evolutionary clusters were detected. An astonishing degree of fine-grained clustering found both between and within our sampling sites underscored the extent to which urban heterogeneity can shape the genetic structure of commensal rodents. Surprisingly, few individuals were recent migrants despite continual global transport, suggesting that recruitment into established populations is limited. Understanding the global population structure of *R. norvegicus* offers novel perspectives on the forces driving the spread of zoonotic disease, and yields greater capacity to develop targeted rat eradication programs.

## INTRODUCTION

The development of agriculture and resultant transition from nomadic to sedentary human societies created new ecological niches for species to evolve commensal or parasitic relationships with humans (Jones, et al. 2013). The phylogeographic history of species living in close association with people often mirrors global patterns of human exploration (Searle, et al. 2009; Gabriel, et al. 2015) and colonization (Matisoo-Smith and Robins 2004; Cucchi, et al. 2005; Suzuki, et al. 2013; Hulme-Beaman, et al. In Press). In particular, commensal rodent distributions have been strongly influenced by the movement of humans around the world. Three rodent species, the house mouse (*Mus musculus*), black rat (*Rattus rattus*), and brown rat (*R. norvegicus*) are the most populous and successful invasive mammals, having colonized most of the global habitats occupied by humans (Long 2003). The least is known about genomic diversity and patterns of colonization in brown rats, including whether a history of commensalism resulted in population divergence, and if so at what spatial scales. Our lack of knowledge of the ecology and evolution of the brown rat is striking given that brown rats are responsible for an estimated $19 billion of damage annually (Pimentel, et al. 2000). Understanding the evolutionary trajectories of brown rats is also a prerequisite to elucidating the processes that resulted in a successful global invasion, including adaptations to a variety of climates and anthropogenic stressors.

We inferred global routes of brown rat expansion, population differentiation, and admixture using a dense, genome-wide nuclear dataset, a first for a commensal rodent (Lack, et al. 2012). A previous mitochondrial study identified the center of origin (Song, et al. 2014) but did not resolve relationships among invasive populations. That work, in combination with fossil distributions (Smith and Xie 2008), suggested that brown rats originated in the colder climates of northern China and Mongolia before expanding across central and western Asia, possibly through human settlements associated with Silk Road trade routes. Based on historical records, brown rats became established in Europe by the 1500s and were introduced to North America by the 1750s (Armitage 1993). Brown rats now occupy nearly every major landmass (outside of polar regions), and human-assisted colonization of islands remains a constant threat to insular fauna (Harper and Bunbury 2015).

Commensalism has given rise to complex demographic and evolutionary scenarios in globally distributed rodents. Although archaeological evidence indicates that commensalism arose long after the emergence of sub-specific lineages in the house mouse in its native range of western Asia (Prager, et al. 1998; Suzuki, et al. 2013), the geographic distribution of *M. m. domesticus* mitochondrial haplotypes reflects transport by humans (Jones, et al. 2010; Suzuki, et al. 2013). *M. m. domesticus* occurred in human settlements along the eastern Mediterranean Basin around 14 kya and rapidly colonized the western Mediterranean and central Europe approximately 3 kya (Cucchi, et al. 2005). Both *M. m. musculus* and *M. m. castaneus* also exhibit regional diversification of mitochondrial lineages due to natural range expansion and spread by human transport (Suzuki, et al. 2013). Human mediated movement has also been implicated in the creation of hybrid zones between subspecies in Scandinavia, China, and New Zealand (Jones, et al. 2010; Jing, et al. 2014; King 2016). Similarly, geographically isolated lineages formed prior to commensalism in the black rat species complex (Aplin, et al. 2011). The spread of agriculture and subsequent trade spurred regional and global range expansion of black rats. Genetic evidence indicates that the global distribution of *R. rattus* Lineage I began with an expansion from the Indian subcontinent into western Asia, followed by separate expansions into Europe and Africa (Tollenaere, et al. 2010; Aplin, et al. 2011). The presence of derived haplotypes also indicates that *R. rattus* Lineage I colonized the Americas, Oceania, and Africa from Europe (Aplin, et al. 2011; Bastos, et al. 2011).

Elucidating global brown rat phylogeographic patterns has several important implications. First, the spread of brown rats may illuminate patterns of human connectivity via trade, or unexpected movement patterns as observed in other commensal rodents (Searle, et al. 2009). Second, rats are hosts to many zoonotic diseases (e.g., *Leptospira interrogans*, Seoul hantavirus, etc.); understanding the distribution of genomic backgrounds may provide insight into differential disease susceptibilities. Additionally, an understanding of contemporary population structure in rats may elucidate source and sink areas for disease transmission. Third, brown rat eradication programs occur in urban areas to decrease disease transmission and on islands where rats prey upon native fauna. A comprehensive understanding of global population structure will allow for better design of eradication efforts, particularly for understanding how to limit new invasions. Thus, our aim was to test biological hypotheses developed from an understanding of the historical narrative of spread using phylogeographic inference. We estimated the number of distinct clusters around the world, the genomic contribution of these clusters within invaded areas, and whether genetic drift and/or post-colonization admixture elicits evolutionary divergence from source populations.

## RESULTS and DISCUSSION

### Evolutionary Clustering

#### Nuclear Genome

Our analyses of 314 rats using 32,127 single nucleotide polymorphisms (SNPs) from ddRAD-Seq identified multiple hierarchical levels of evolutionary clustering (K). Principal component analysis (PCA) distinguished two clusters along the first principal component (PC), an Asian cluster that extended to western North America, and a non-Asian cluster found in Europe, Africa, the Americas, and New Zealand (Fig. 1). Higher dimension PCA axes distinguished subclusters (Fig. S2), then individual sampling sites; in total 58 axes of variation were significant using Tracy-Widom statistics (20 and 37 axes were significant for PCAs with only Asian or non-Asian samples respectively). Using the model-based clustering program ADMIXTURE, the Asian and non-Asian clusters divided into five and eight subclusters, respectively (Fig. 2, 3, S3-S5). Higher numbers of clusters (K=18, 20, and 26) were also supported by ADMIXTURE (Fig. S3A, S4), distinguishing ever finer spatial scales (from subcontinents to cities).

**Fig. 1.**
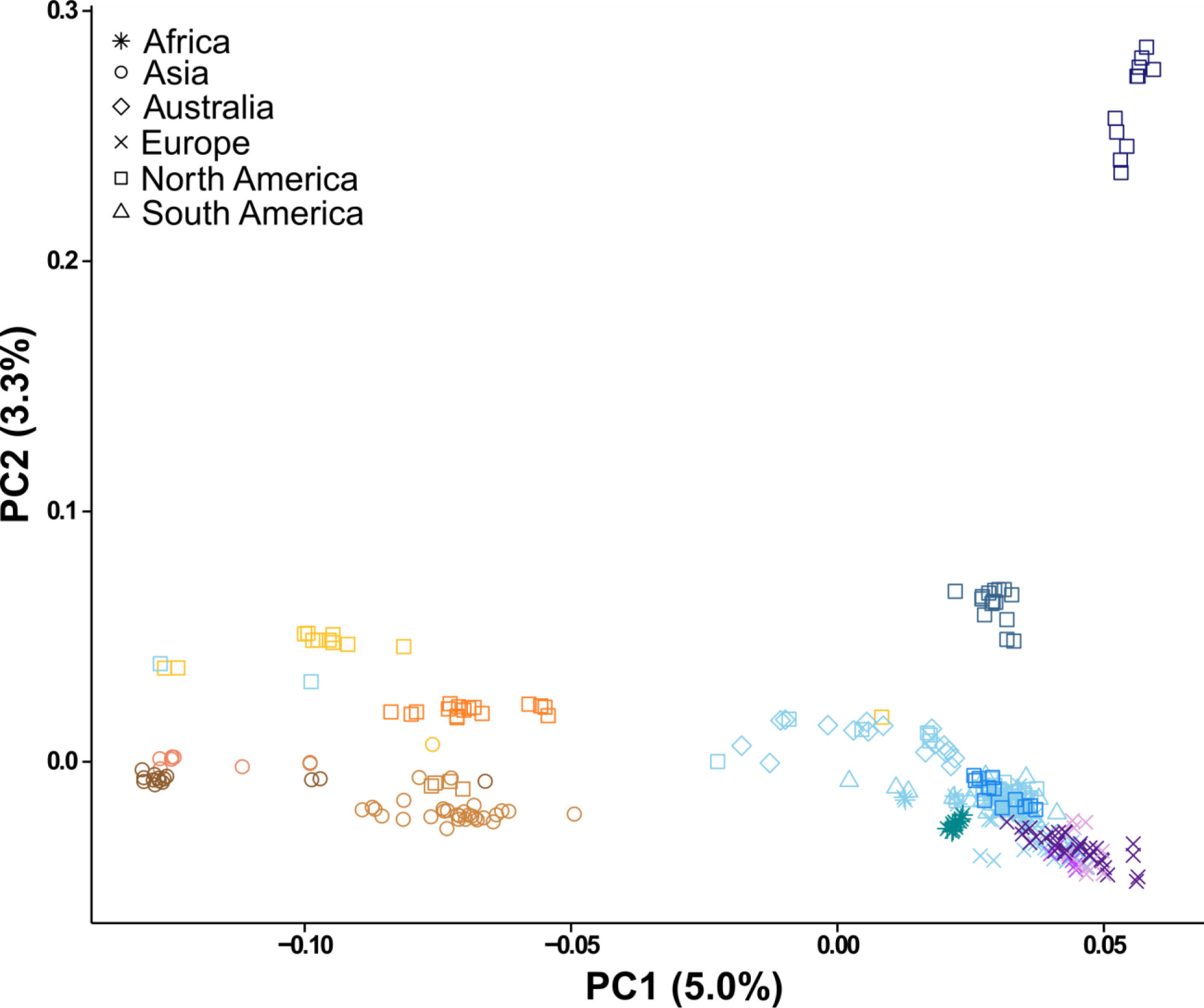
Principal components analysis using 32k nuclear SNPs for worldwide *Rattus norvegicus* samples for the first two principal components. Continents are designated by shape (Asia: circles; Europe: X; Africa: star; North America: square; South America: triangle; New Zealand: diamond) with substructured populations designated by color for the 13 clusters inferred using model-based ancestry analyses (Figs. 2, S4).

The subclusters in the Asian cluster reflect underlying geography and hierarchical differentiation (Fig. S3B). The predominant four clusters reflected differentiation between: China, Southeast (SE) Asia, the Aleutian Archipelago, and Western North America (Fig. S6, S7). Within the SE Asia cluster, further subdivision was observed for the Philippines and Thailand (Fig. 2, S7). Within the Aleutian Archipelago cluster, samples from the city of Sitka (in the Alexander Archipelago) formed a subcluster. Rats from the Russian city of Sakhalinskaya Oblast and four rats aboard the Bangun Perkasa ship each formed a subcluster (Fig. S7). The Bangun Perkasa was a nationless vessel seized in the Pacific Ocean by the US government in 2012 for illegal fishing. Our analyses identified that the rats aboard were of SE Asian origin and likely represented a city in that region, probably one bordering the South China Sea, at which the ship originated or docked.

**Fig. 2.**
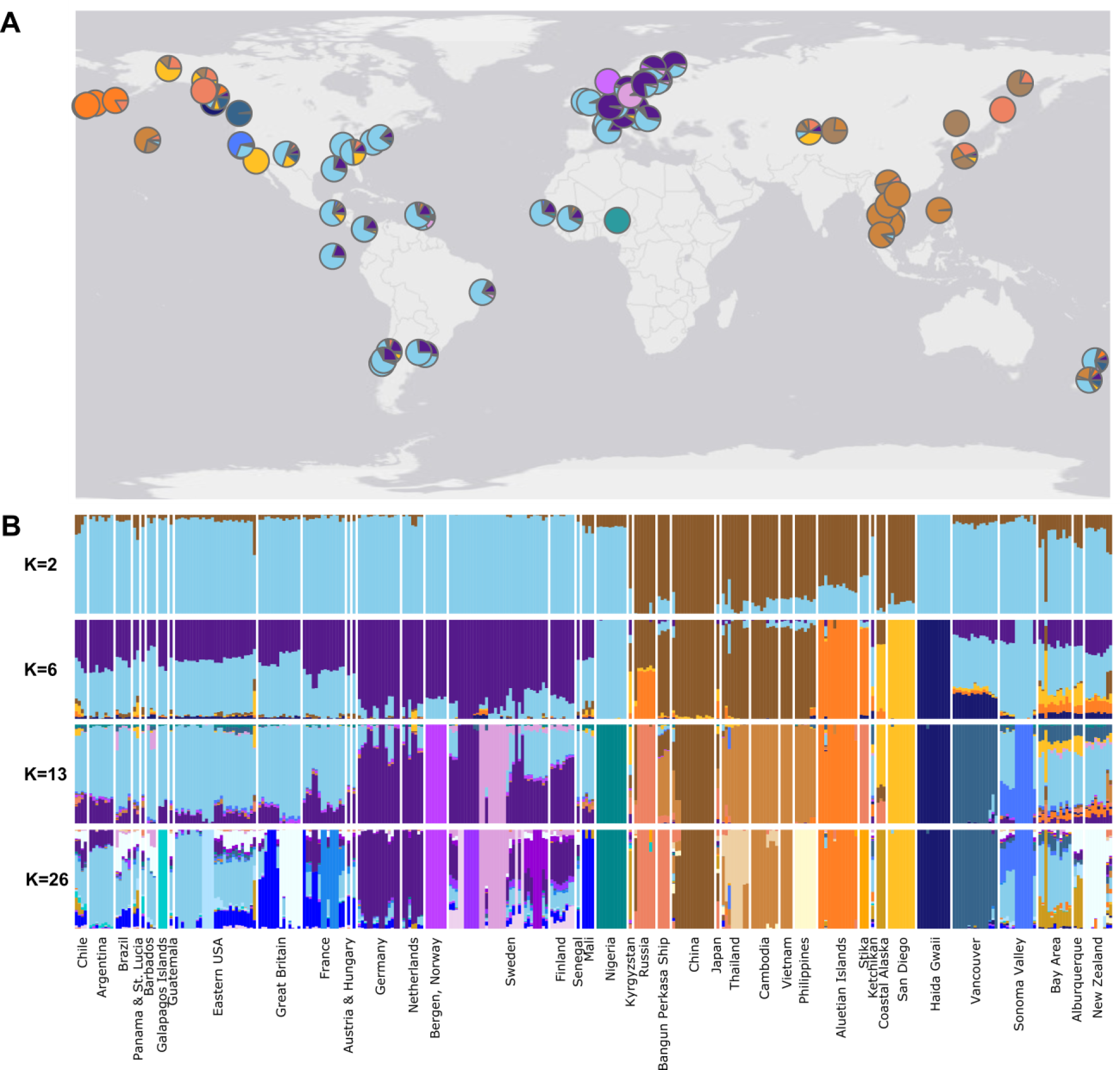
(A) Map of brown rat sampling locations with average proportion of ancestry per site inferred using 32k nuclear SNPs. Ancestry was based on ADMIXTURE estimates from 13 clusters (China: brown; SE Asia: light brown; Russia: pink; Aleutian Archipelago: orange; western North America: gold; W Euro: light blue; N Euro: purple; Kano: turquoise; Sonoma Valley: medium blue; Haida Gwaii: dark blue; Vancouver: cerulean; Bergen: medium purple; Malmo: light purple). (B) Ancestry proportions from ADMIXTURE for 314 samples at two, six, 13, and 26 clusters.

We detected greater hierarchical differentiation in the non-Asian cluster (Fig. S3C). At K=3 we observed divergence between the Western Europe (W Euro) and Northern Europe (N Euro) clusters (Fig. S9). The W Euro cluster contained rats from Europe (Great Britain, France, Austria, and Hungary), Central and South America (Argentina, Brazil, Chile, Galapagos Islands, Honduras, Guatemala, Panama), the Caribbean (Barbados, Saint Lucia), North America (eastern, central, and western USA), New Zealand, and Africa (Senegal and Mali); and the N Euro cluster included Norway, Sweden, Finland, Germany, and the Netherlands (Fig. 2, S4, S8, S9). Within these broad geographic regions, many subclusters were identified by ADMIXTURE that likely resulted from either intense founder effects, isolation resulting in genetic drift, the inclusion of second and third order relatives in the dataset, or a combination of these factors. In the global analysis, four clusters were nested within W Euro (the island of Haida Gwaii, Canada; Vancouver, Canada; Kano, Nigeria; and Sonoma County in the western USA) and two within N Euro (Bergen, Norway; Malmo, Sweden). We identified additional well-supported subclusters within the non-Asian cluster at K=12, 15, and 17 that represented individual cities (Fig. S9).

Our analysis using FINESTRUCTURE identified 101 clusters (Fig. 3). Of the 39 cities where more than one individual was sampled, 19 cities supported multiple clusters indicating genetic differentiation within cities. As GPS coordinates were not collected, we cannot hypothesize if these clusters represent distinct populations or were artefacts of sampling relatives, despite removal of individuals with relatedness coefficients greater than 0.20, although the FINESTRUCTURE algorithm should be robust to relatedness when identifying clusters. The Asian and N Euro sampling sites individually had higher coancestry coefficients between locations (Fig. 3) which supported the hierarchical clustering observed using ADMIXTURE.

**Fig. 3.**
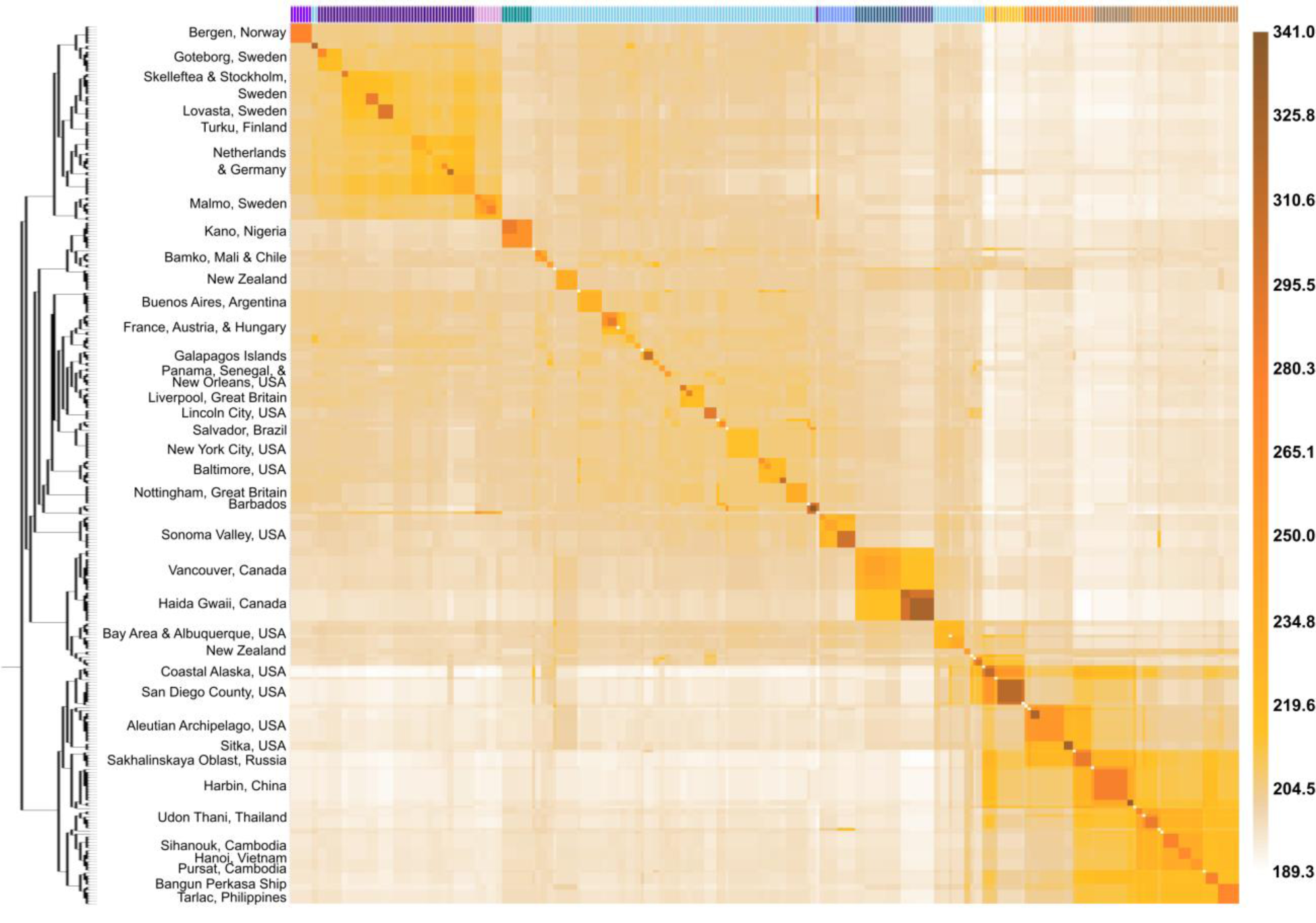
Coancestry heat map of brown rats, where light and dark brown, respectively denote lower and higher coancestry. The 101 populations identified by FINESTRUCTURE appear along the diagonal. A bifurcating tree and select sampling locations are shown on the left, and assignment to one of the 13 clusters from Fig. 2 shown on top.

#### Mitochondrial Genome

We identified 10 clades within a network-based analysis of 103 mitochondrial haplotypes (Fig. 4, Tables S5, S6). Many of the clades had spatial structure concordant with the nuclear genome results (Fig. 2A). We observed clade 1 in China, Russia, and western North America. Additionally, clades 6 and 9 contained a single haplotype only observed in China. We interpret the diversity of clades within northern China as representative of geographic structure in the ancestral range prior to movement of rats by humans (Fig. 4, Table S6). In SE Asia we observed clades 2 (aboard the Bangun Perkasa), 3 (Philippines), and 5 (Cambodia, Thailand, and Vietnam). Clade 4 was found in western North America. European samples comprised three divergent clades (3, 8, and 10). Clade 8 was observed across Europe, western North America, and South America; this clade shared ancestry with clade 7 which was observed in Russia and Thailand (Fig. 4).

**Fig. 4.**
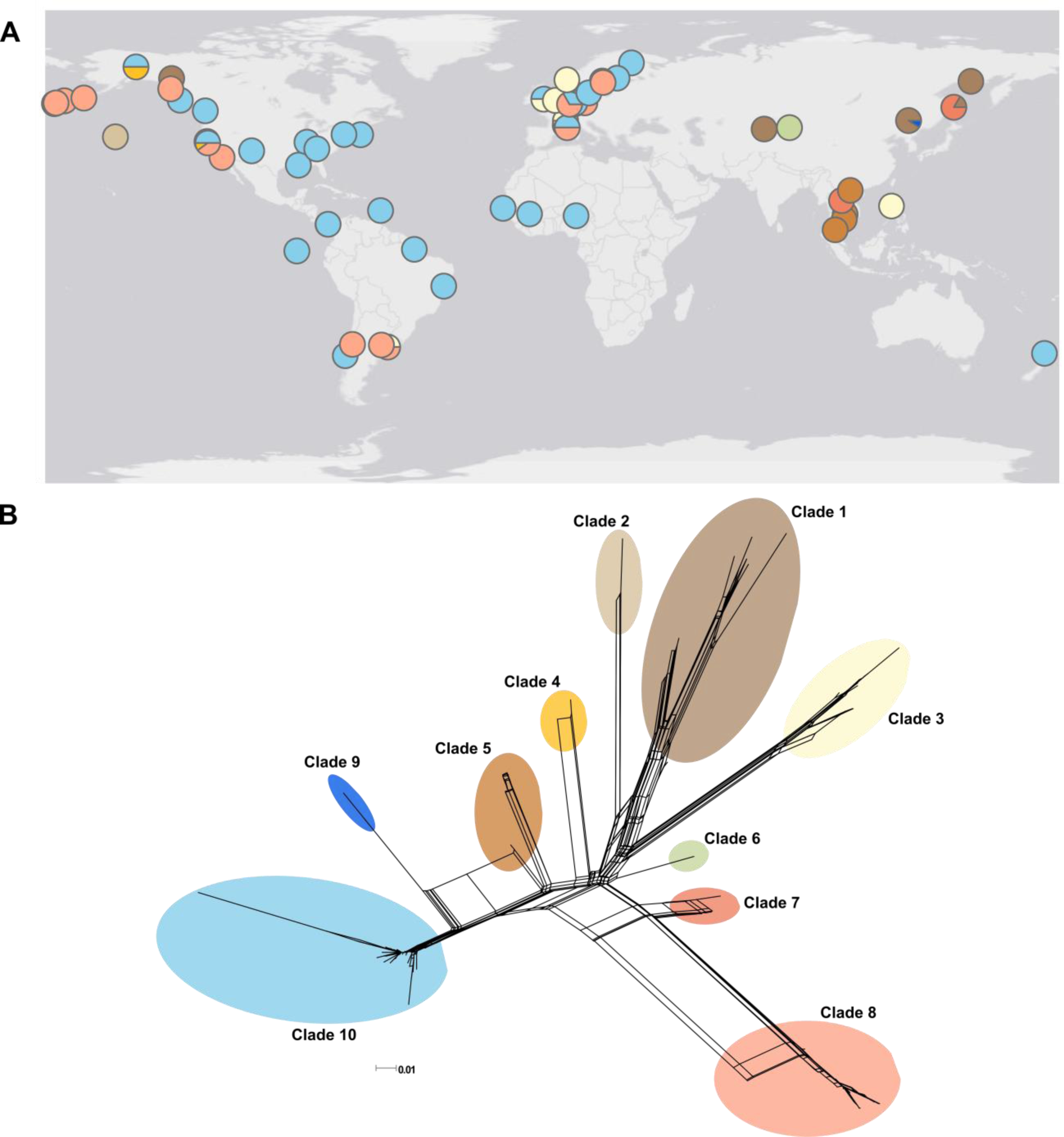
(A) Map of the proportion of mitochondrial clades at each sampling site for 144 individuals and (B) SNP haplotype network with 103 haplotypes in 10 clades (clade 1: brown; 2: beige; 3: pale yellow; 4: gold; 5: light brown; 6: pale green; 7: pink; 8: light pink; 9: dark blue; 10: light blue).

### Range Expansion

We thinned our dataset to the sampling site with the largest sample size within each of the 13 clusters supported by ADMIXTURE and analyzed the data using TREEMIX (Fig. S10). We observed divergence within Asia first, followed by the two independent expansions into western North America. Drift along the backbone of the non-Asian cluster was limited, indicating rapid expansion of rats into Africa, Europe, and the Americas (Fig. S10). Both the population tree topology and PCA (Figs. 1, S2, S10) indicated that range expansion occurred in three directions, where one southward and two eastward expansions comprised Asian ancestry, and the westward expansion produced the non-Asian cluster.

#### Ancestral Range

In eastern China, the nuclear genome assigned strongly to a single cluster while mitochondrial diversity encompassed two divergent clades, where samples from western China assigned to both the Chinese and SE Asian clusters and represented a third mitochondrial clade. This result suggests substructure within the ancestral range, although the samples from northeastern China may not be representative of the ancestral range but instead of an isolated, divergent population that has retained high genetic diversity (Tables S4, S6).

#### Southern Expansion into SE Asia

A southward range expansion into SE Asia was supported by the population tree topology, higher heterozygosity, low nuclear *F*_ST_ with China, and elevated coancestry coefficients between populations in SE Asia, China, and Russia (Fig. 3, Tables S3, S4). Given evidence for an early southward expansion (Fig. S10), we hypothesize that the founding of SE Asia was accompanied by a weak bottleneck resulting in relatively low loss of genetic diversity. However, following founding regional diversification occurred as we observed substructure in both the nuclear and mitochondrial genomes (Fig. 2, 4, S7).

#### Two Independent Eastward Expansions

We observed population divergence along the first eastward expansion from eastern Russia into the Aleutian Archipelago based on PCA (Fig. S6). Both the population tree topology and PCA indicate that a second eastward expansion progressed from Asia to western North America (Fig. S6, S10). While the Western North America cluster was observed in both northern and southern Pacific coast localities (Fig. S5A), we cannot extrapolate that this cluster represents the entirety of the coastline. Specifically, Sitka, Ketchikan, Vancouver, and the Bay Area are all located between the Alaskan cities and San Diego County that comprise the Western North America cluster. Further, the timing of these expansions is an open question. While the population tree indicated divergence of these two expansions prior to divergence of the non-Asian cluster, the historical record attributes brown rats in the Aleutian Archipelago to Russian fur traders in the 1780s (Black 1983), which is not consistent with rats entering Europe in the 1500s (Armitage 1993). Thus, evidence of early divergence may be a consequence of unsampled Asian populations sharing ancestry with the Aleutian Archipelago and Western North America clusters.

#### Westward Range Expansion into Europe

The low drift along the backbone of the population tree for the non-Asian cluster is indicative of rapid westward expansion (Fig. S10). Limited inferences could be drawn about western Asia and the Middle East because of sampling constraints, but we hypothesize that the region was colonized by the range expansion of the non-Asian cluster. We observed three mitochondrial clades in Europe, where clade 3 shared ancestry with SE Asia and clade 8 shared ancestry with eastern Russia, while clade 10 is a European derived clade (Fig. 3, Table S6). Thus, Europe may have been independently colonized three times, although the routes remain an open question. We hypothesize that clade 10 arrived overland around the Mediterranean Sea, similar to black rats (Aplin, et al. 2011). We hypothesize that following the independent colonizations, the genetic backgrounds admixed prior to divergence between the N Euro and W Euro clusters given the low nuclear *F*_ST_ (Table S4).

Notably, we detected genetic differentiation of Bergen, Norway and Malmo, Sweden within the N Euro cluster (Fig. 2). This pattern suggests drift following either a strong founder effect or population isolation and limited gene flow. Isolation is likely driving the pattern observed in Bergen, which is separated from eastern Norway by mountains that are thought to limit movement of commensal rodents (Jones, et al. 2010).

### Range Expansion of Rats by Europeans

We detected a fifth range expansion that can be attributed to transport by western European imperial powers (1600s-1800s) to former colonial territories (Fig. 2, 3, S4, S9). For example, we observed high proportions of W Euro ancestry in samples from the North and South Islands of New Zealand, which is consistent with the introduction of brown rats by British colonists, as has also been inferred for black rats (Aplin, et al. 2011) and domestic cats in Australia (Spencer, et al. 2016). We observed admixture on both islands (Fig. 3) although nuclear ancestry proportions differed between the islands with higher proportions of N Euro and Vancouver ancestry on the North Island. The South Island had higher SE Asia and Western North American ancestry (Fig. 2, 3, S4); these ancestry components may be attributed to the seal skin trade with southern China by sealers from the USA (King 2016).

The samples from Nigeria and Mali formed a sister clade in FINESTRUCTURE, which likely reflects a shared history as French colonies, although Senegal fell outside of the clade (Fig. 3). Mali had elevated W Euro ancestry compared to Nigeria which may be a consequence of multiple introductions from European sources. South American countries exhibited a paraphyletic FINESTRUCTURE topology that is suggestive of colonization from multiple locations. This result was also supported by the presence of all three mitochondrial clades found in Europe (Fig. 4A). Further sampling from Portugal and Spain would better resolve the origins of Brazilian populations and clarify relationships of former colonies elsewhere in the world.

The complex distribution of clusters in North America is suggestive of a dynamic colonization history, including independent introductions on both the Atlantic and Pacific coasts (Fig. 2). We detected mtDNA haplotypes of European ancestry in the eastern and central USA, whereas the Pacific seaboard harbors high mtDNA haplotype diversity from European and Asian clades (Fig. 4). These results are consistent with prior observations of four high-frequency mtDNA haplotypes across Alaska and the continental USA, of which three were observed in east Asia and one in Europe (Lack, et al. 2013). Along the Pacific coast, cities with both Asian and non-Asian nuclear ancestry were observed (Fig. 2), which parallels the pattern observed in black rats (Aplin, et al. 2011). Given the bicoastal introductions, it is unsurprising to observe admixture in North American cities such as the San Francisco Bay Area and Albuquerque, where each has elevated coancestry coefficients with Asian and non-Asian clusters (Fig. 3). We also observed limited eastward dispersal of Asian genotypes, although other work has found evidence of greater inland penetration (Lack, et al. 2013).

Rats from Haida Gwaii off the coast of British Columbia, Canada, were consistently recovered as a separate cluster in ADMIXTURE, and had high coancestry coefficients and *F*_ST_ with other populations (Fig. 3, Table S4), indicating substantial genetic drift following colonization. Rats were introduced to Haida Gwaii in the late 1700s via Spanish and/or British mariners, and have been subject to recent, intensive eradication efforts that may have heightened genetic drift (Hobson, et al. 1999).

### Intra-urban Population Structure of Brown Rats

Brown rats exhibit population structure over a remarkably fine-grained spatial scale (Fig. 3); specifically, rat population structure exists at the scale of both cities and neighborhoods. We found evidence of heterogeneity among cities as some appear to support one population while others support multiple populations. For example, we detected a single population across multiple neighborhoods in Manhattan (NYC, USA), whereas four genetic clusters (Fig. 3) were observed in a neighborhood in Salvador, Brazil, a result that confirmed previous microsatellite based analyses (Kajdacsi, et al. 2013). Although denser sampling will be needed to confirm whether these groups represent distinct populations or reflect oversampling of intra-city pockets of highly related individuals, intra-city clustering likely represents substructure considering the global design of our SNP dataset. Observations of highly variable intra-city structure suggest the following three scenarios: first, effective population size rapidly increases after invasion, possibly driven by high urban resource levels, and thus genetic drift may have a relatively weak effect on population differentiation. Second, new immigrants that arrive after initial invasion and establishment of rats in a city may be limited in their capacity to either establish new colonies or join existing colonies (Calhoun 1962), thereby limiting ongoing gene flow from other areas due to competitive exclusion (Waters 2011). Gene flow into colonies may also be sex-biased as females were recruited more readily than males in a two-year behavioral study of brown rats (Calhoun 1962). We did observe gene flow in our dataset, including an individual matching Coastal Alaska into the Bay Area and an individual with high Sonoma Valley ancestry in Thailand (Fig. 2B), thus migration due to contemporary human-assisted movement is possible and ongoing. However, given increasing connectivity due to trade and continual movement of invasive species (Banks, et al. 2015), we expected greater variability in ancestry proportions within cities than observed (Fig. S4). Third, cityscapes vary in their connectivity where some cities contain strong physical and/or environmental barriers facilitating differentiation and others do not. Identifying commonalities and differences among cityscapes with one or multiple rat populations should be a goal for understanding how rats interact with their environment, particularly in relation to the effect of landscape connectivity for pest and disease control efforts.

### Significance

#### Understanding the Spread of Zoonotic Pathogens

Understanding the global population structure of brown rats offers novel perspectives on the forces driving the spread of zoonotic disease. Our inference that competitive exclusion may limit entry into established populations helps explain why zoonotic pathogens do not always exhibit the same spatial distribution as rat hosts as well as the patchy distribution of presumably ubiquitous pathogens within and between cities (Himsworth, et al. 2013). While within-colony transmission of disease and natal dispersal between colonies are important factors related to the prevalence of zoonotic disease, our results also suggest that contemporary human-aided transport of infected rats does not contribute to the global spread of pathogens, as we would expect higher variability of ancestry proportions within cities if rats were successfully migrating between cities. Additionally, our results indicate that rats with different genomic backgrounds may have variable susceptibilities to pathogens, though differential susceptibility likely depends on concordance between the geographic origins of pathogens and rats. While this idea needs pathogen specific testing, it could have substantial implications for global disease transmission.

#### Rat Eradication Programs for Species Conservation

Eradication of invasive *Rattus* species on islands and in ecosystems with high biodiversity is a priority for conservation of at-risk species, as rats outcompete or kill native fauna. It remains challenging to gauge the success of eradication programs, because it is difficult to distinguish between post-intervention survival and reproduction as opposed to recolonization by new immigrants (Piertney, et al. 2016). Understanding fine-scale population genetic structure using dense nuclear marker sets (Robins, et al. 2016), as in this study, would allow managers to more clearly assess outcomes and next steps following an eradication campaign. For example, genomic analyses could illustrate that an area has been recolonized by immigration from specific source populations, thereby allowing managers to shift efforts towards biosecurity to reduce the likelihood of establishment by limiting the influx of potential immigrants.

## MATERIALS and METHODS

We obtained rat tissue samples from field-trapped specimens, museum or institute collections, and wildlife markets (Tables S1, S2). As GPS coordinates for individuals were not always available, the sampling location was recorded as either the city, nearest town, or island where rats were collected.

### DNA Extraction, RAD sequencing, and SNP calling

We extracted DNA following the manufacturer’s protocols using Qiagen DNeasy kits (Valencia, CA). We prepared double digest restriction-site associated DNA sequencing (ddRAD-Seq) libraries with 500-1000ng of genomic DNA from each sample and one negative control made up of water. Briefly, samples were digested with SphI and MluCI before ligation of unique barcoded adapters. We pooled 48 barcoded samples each in 10 libraries at equimolar concentrations. We then selected fragments from 340-412 bp (target = 376 bp) using a Pippin Prep (Sage Science, Beverly, MA). The size-selected pools were PCR-amplified for 10-12 cycles using Phusion PCR reagents (New England Biolabs, Ipswich, MA) and primers that added an Illumina multiplexing read index. Final libraries were checked for concentration and fragment size on a BioAnalyzer (Agilent Technologies, Santa Clara, CA), then sequenced (2 × 125bp paired-end) at the New York Genome Center across five lanes of an Illumina HiSeq 2500.

We demultiplexed the raw reads using the process_radtags script in STACKS v1.35 (Catchen, et al. 2013), then aligned reads for each individual to the *Rattus norvegicus* reference genome (Rnor_6.0) (Gibbs, et al. 2004) using Bowtie v2.2.6 (Langmead and Salzberg 2012) with default parameters. To assess the number of mismatches allowed between stacks and the minimum depth of coverage for each stack when building RADtags (-n and -m flags respectively) in STACKS, we processed two samples under a number of scenarios and compared the number of RADtags that formed as de novo loci versus those that mapped to the reference *R. norvegicus* genome. We first assessed the M parameter by holding m constant at three while varying M between two and five in the ustacks program. We observed a decrease in the undermerged RADs with increasing values of M; we selected M = 4 for both the final RAD processing and as the constant level when we allowed m to vary between two and five. We selected m = 3 to balance between removing real loci and stacks that erroneously mapped to the reference genome. In the cstacks program we assessed the number of allowed mismatches between tags (n) from zero to two. We observed little difference for this parameter between our test values and decided to use n = 2 as a conservative measure.

We initially built the STACKS catalog with all of the reference-aligned samples (n = 447) using the ref_map pipeline. Following processing, we filtered for the following: biallelic SNPs, a minor allele frequency (MAF) greater than or equal to 0.05, SNPs genotyped in 80% of samples, and only one SNP per RADtag (STACKS flag --write_single_snp); additionally, SNPs that mapped to either the Y chromosome or mitochondrial genome were removed. This dataset had 37,730 SNPs. Following sample collection and genotyping, we were informed that *R. rattus* samples had been collected in Mali; we capitalized on this by confirming the species identification for each sample using principal components analysis (PCA) in EIGENSOFT v5.0.2 (Patterson, et al. 2006; Price, et al. 2006), and ADMIXTURE v1.23 (Alexander, et al. 2009) for two clusters. We identified 33 *R. rattus* and 414 *R. norvegicus* samples (Fig. S1).

We reran ref_map using only the confirmed *R. norvegicus* samples, and filtered similarly as described above plus an additional filter to remove individuals with greater than 60% missing data. To add genotypes from 11 of the *R. norvegicus* samples collected in Harbin, China (European Nucleotide Archive ERP001276) (Deinum, et al. 2015), we mapped reads to the Rnor_6.0 genome using SAMTOOLS v1.2 (Li, et al. 2009) then extracted the SNP dataset using mpileup with a position list. We removed related individuals within, but not between, sampling sites by assessing relatedness in KING v1.4 (Manichaikul, et al. 2010). For each pair of individuals with relatedness estimators greater than 0.2, one individual was removed from the analysis (n = 22). Subsequently, we randomly thinned 14 samples from Vancouver, Canada as preliminary analyses indicated oversampling. Thus the final nuclear *R. norvegicus* dataset contained 32,127 SNPs genotyped in 314 individuals (Table S1).

From the initial processing in STACKS, we extracted the SNPs that mapped to the mitochondrial genome to produce a second dataset with 115 SNPs (see Table S5 for base pair positions within the *R. norvegicus* reference mitochondrial genome, GenBank accession AY172581.1). We extracted the same positions from the mitochondrial genomes of samples from Harbin, China. We allowed up to 35% missing data per individual and identified 103 haplotypes using COLLAPSE v1.2 (Posada 2004) in 144 individuals. We built a haplotype network using SPLITSTREE v4.13.1 (Huson and Bryant 2006) and identified the haplotypes grouped into 10 clades (Table S6).

### Population Genomic Analyses

To describe population structure, we ran ADMIXTURE (Alexander, et al. 2009) at each cluster from 1 to 40. Given known effects of sampling bias on clustering analyses, we repeated this analysis with a subset of the data where four or five samples from each city were randomly selected (n = 158). The results supported K=14 clusters which supported the analysis of our full dataset. We also subdivided the full dataset into the Asian and non-Asian clusters and reran ADMIXTURE at each cluster from 1 to 25. We used the CV error values to identify the best-supported clustering patterns across the range. Using the same datasets (full, Asian, and non-Asian), we ran PCA in EIGENSOFT (Patterson, et al. 2012) and identified significant PCs using Tracy-Widom statistics.

We also estimated evolutionary clusters using FINESTRUCTURE v2.0.7 (Lawson, et al. 2012) which elucidates the finest grained clusters by accounting for linkage disequilibrium and allows detailed admixture inference based upon the pairwise coancestry coefficients. We limited this analysis to the 20 autosomes (31,489 SNPs), removing SNPs on unassembled scaffolds in the dataset. Data for each chromosome were phased and imputed using fastPHASE v1.2 (Scheet and Stephens 2006). Initial analyses using the linked model indicated our data were effectively unlinked (c-factor 0.0104); therefore, we ran the unlinked model. We used default settings except for the following parameters: 25% of the data were used for initial EM estimation; 750,000 iterations of the MCMC were run (375,000 of which were burnin) with 1,000 samples retained, 20,000 tree comparisons, and 500,000 steps of the tree maximization were run. We viewed MCMC trace files to confirm stability of all parameters.

To understand patterns of population divergence, we ran TREEMIX v1.12 (Pickrell and Pritchard 2012). As the *R. rattus* data (see Supplemental Methods) were mapped to the *R. norvegicus* genome, we extracted SNPs at the same genomic positions for 31 black rats (we removed two samples showing admixture; Fig. S1) with SAMTOOLS (Li, et al. 2009) mpileup function using a position list. We selected the sampling location with the largest sample size from each of the well supported clusters at K=13 (Fig. 2, S4), plus the *R. rattus* samples for the outgroup (which were not subdivided due to lack of population structure, Fig. S11). We added migration edges to the population tree sequentially by fixing the population tree to the tree with n-1 migration edges, where blocks of 1,000 SNPs and the sample size correction were enabled. We assessed both the proportion of variance (Fig. S12A) and the residuals of the population tree (Fig. S12B) and chose the model with three migration edges. We decided to thin the sampling areas due to uneven sampling between the broad Asian and non-Asian clusters; both factors should affect the variance in the model, thus we presented a potentially underfit versus overfit model. We ran *f*_3_ tests within TREEMIX and observed no significant relationships, likely due to highly complex admixture patterns (Patterson, et al. 2012).

For the nuclear dataset, we calculated expected heterozygosity (H_E_) and *F*_IS_ within each of the 13 clusters using ARLEQUIN v3.5.1.3 (Excoffier and Lischer 2010), and pairwise *F*_ST_ using VCFTOOLS v0.1.13 and the Weir and Cockerham estimator (Weir and Cockerham 1984; Danecek, et al. 2011). For the mitochondrial dataset, we calculated pairwise *F*_ST_ between the clusters identified in the nuclear dataset in ARLEQUIN.

## ACKNOWLEDGEMENTS

This work was funded by National Science Foundation grants DEB 1457523 and DBI 1531639, and a Fordham University faculty research grant, to JM-S. We thank Kaitlin Abrams and Ian Hays for assisting with lab work. We thank Annette Backhans, Francois Catzeflis, Gauthier Dobigny, Carol Esson, Tim Giles, Gregory Glass, Sabra Klein, Mare Lõhmus, Patrick McClure, Frank van de Goot, Jordan Reed and his Mongrel Hoard, Richard Reynolds and the Ryders Alley Trencher-fed Society (R.A.T.S), Thomas Persson Vinnersten and colleagues at Anticimex, and partners of the Network Rodent-Borne Pathogens for collecting and providing rat samples. The mammal collections at the University of Alaska Museum of the North, Angelo State University, Berkeley Museum of Comparative Zoology, the Burke Museum at University of Washington, the Museum of Southwestern Biology, and the Museum of Texas Tech University also graciously provided tissue samples.

